# Ranging and territoriality in African Apes

**DOI:** 10.64898/2026.11.03.763879

**Authors:** Martin Surbeck, Rebecca F. Supple, Anthony P. Massaro, Deus C. Mjungu, Martha M. Robbins, Michael L. Wilson

## Abstract

Intergroup relations vary strikingly among African apes, with only chimpanzees showing intense, often lethal territoriality. The economic defendability hypothesis proposes that such territorial behaviour is favored where home ranges can be economically defended, as indexed by the ratio of daily travel distance to home range diameter. To test this idea, we compiled and analyzed comparable data on yearly home range size, daily travel distance and defendability indices from one population of each nonhuman African ape species and integrated these with comparable published data. We find that bonobos had substantially larger home ranges and longer daily travel distances than chimpanzees and gorillas, contradicting a prevailing assumption that bonobos experience higher or more reliable food abundance. Defendability indices were higher in chimpanzees and bonobos than in mountain gorillas, but did not differ significantly between chimpanzees, bonobos, and western gorillas. Overall, our results indicate that economic defendability alone cannot explain interspecific variation in territoriality and that our current understanding of ecological differences between the species of the genus Pan have to be revised. Possible additional factors shaping space use of neighboring groups include mate competition within groups.

## INTRODUCTION

Intergroup behaviour likely played a major role in shaping the evolution of human behavior, but vigorous debate continues over the relative importance of peaceful versus violent interactions between groups in human prehistory (Fry, 2013; Glowacki, 2024; Majolo, 2019; Meijer, 2024). Studies of our living primate relatives offer insights into the selective pressures and processes shaping different forms of intergroup interactions (Pisor & Surbeck, 2019) and can inform conceptual models of human evolution. The African apes, the clade to which humans belong, vary considerably in their patterns of intra- and intergroup relations (Furuichi, 2020). Consequently, researchers interested in the evolution of war and peace in humans have attempted to explain the striking contrasts in intergroup behaviour between our closest living relatives: chimpanzees (*Pan troglodytes*) and bonobos (*Pan paniscus*) (Glowacki, 2024; Meijer, 2024; R. Wrangham, 1999). Chimpanzees are territorial and with the exception of potential female immigrants, are hostile towards and sometimes kill members of other groups (Wilson et al., 2014). Intergroup relations among bonobos can be hostile, but prolonged peaceful intergroup mingling also occurs (Lucchesi, Cheng, Janmaat, et al., 2020; Tokuyama et al., 2019).

Efforts to explain these strikingly different intergroup relations of chimpanzees and bonobos have focused on ecological differences between the two species. Social factors, such as patterns of philopatry and dispersal, are known to shape intergroup behavior across primates more broadly (Willems et al., 2013; Willems & van Schaik, 2015), but are unlikely to account for the contrast between chimpanzees and bonobos, given that both species share a similar social system characterized by male philopatry and female dispersal. However, bonobos are considered to have more cohesive parties, making them less vulnerable to lethal raiding (R. W. Wrangham, 1999). The more cohesive parties of bonobos are, in turn, proposed to result from access to more abundant or reliable food supplies (Malenky, 1990; White & Wrangham, 1988; R. Wrangham, 2023). In contrast, Pusey (Pusey, 2022, p. 202) noted that bonobos appear to have larger home ranges than chimpanzees, suggesting that they have home ranges that are not economical to defend, such that aggressive territorial behavior is not likely to provide fitness benefits for bonobos. While economic defendability unlikely provides a complete explanation for all the observed differences in intergroup behavior, it can help explain some of the contrasting patterns.

The two gorilla species, western gorillas (*Gorilla gorilla*) and eastern gorillas (*Gorilla beringei*), with its longest studied subspecies, mountain gorillas (*Gorilla beringei beringei*), provide additional valuable comparative data, as they occupy many of the same habitats as chimpanzees and bonobos and share a very similar ecological niche. In both western gorillas and mountain gorillas, encounters between groups often include aggression, including conspicuous displays by adult males, and sometimes escalate into direct physical confrontations, including infanticidal attacks (A. M. Robbins et al., 2013) and, in rare cases, killing of adults (Rosenbaum et al., 2016). Nonetheless, peaceful interactions between groups are also regularly observed (Forcina et al., 2019; M. M. Robbins & Sawyer, 2007; Rosenbaum et al., 2016).

Differences in ecological variables that shape travel patterns and space use (Koenig, 2002) are considered relevant to intergroup behavior, as they determine whether defending a home range is economically feasible in the first place (Mitani & Rodman, 1979). A home range is the area where an animal (or social group) normally lives (Brown & Orians, 1970). A territory is a home range that is (1) relatively fixed in area, (2) actively defended against intruders, and (3) exclusively used by the resident(s) (Brown & Orians, 1970). Among the nonhuman African apes (hereafter, “African apes”), gorillas are generally not considered territorial and have extensively overlapping home ranges (Caillaud et al., 2014; Judson et al., 2024; Seiler et al., 2017). However, both gorilla species exhibit some elements of territorial behavior, with mountain gorillas having exclusive core areas (Morrison, Hirwa, et al., 2020; Seiler et al., 2017) and western gorillas being more hostile to outsiders in the core of their home range than in the periphery (Morrison, Dunn, et al., 2020). At long-term chimpanzee study sites with multiple neighboring communities there is little home range overlap between neighboring communities and mainly males have been described to defend group territories and conduct boundary patrols (Lemoine et al., 2020; Wilson & Wrangham, 2003). The home ranges of bonobos overlap considerably, and while bonobo groups do compete over access to food resources, there is little evidence that bonobos actively defended their home ranges (Samuni et al., 2022; Tokuyama et al., 2019).

According to the *economic defendability hypothesis*, territorial behaviour can occur when a home range is economically defendable, in which case group members may try to exclude outsiders. Economic defendability can be measured by an index (D), which consists of the ratio of the mean daily travel distance to the annual home range diameter (Mitani & Rodman, 1979). Mitani and Rodman (Mitani & Rodman, 1979) found that all populations that were described as territorial had D ≥ 1, whereas many populations described as non-territorial exhibited lower values. Generally across primates, larger home ranges are more difficult to patrol (Pearce et al., 2013) and Pusey (Pusey, 2022) suggests that differences in the *size of home ranges* between chimpanzees and bonobos may underlie the observed variation in the intensity of intergroup competition, with the larger home ranges of bonobos being less easily defended than those of chimpanzees. However, formal comparisons of home range sizes between the two species and the ones of gorillas have not yet been conducted.

*A priori* several ecological factors generate distinct predictions about home range size variation among African apes. First, across primates, energetic requirements scale with body mass, such that individuals of larger-bodied species must forage across larger areas to meet their metabolic needs. Given that gorillas are substantially larger-bodied than *Pan* species, this allometric relationship predicts larger home ranges in gorillas (Prediction 1a: Home range size: MG/WG>C/B). Second, an influential ecological hypothesis posits that species with greater dependence on fruit require larger home ranges to meet their energetic needs, since fruit resources tend to be more spatially and temporally dispersed than foliage (Clutton-Brock & Harvey, 1977; Milton & May, 1976). This relationship, however, does not appear to be universal, as highly folivorous taxa such as *Rhinopithecus* can exhibit some of the largest home ranges reported among primates (Kirkpatrick & Grueter, 2010), indicating that low frugivory does not necessarily correspond to small ranges and that other ecological variables such as habitat productivity can be important (Kamilar & Baden, 2014; Tucker et al., 2014; Willems et al., 2013). Finally, larger social groups are generally expected to experience greater within-group feeding competition and consequently require larger ranging areas to meet the collective nutritional needs of their members.

Among the African apes, gorillas eat the least fruit, though the proportion of fruit in gorilla diets varies considerably: less than 1% of food intake for Virunga mountain gorillas (Watts, 1984), about 15% of feeding time in Bwindi mountain gorillas (Ostrofsky & Robbins, 2020), and roughly 30% of the feeding time for western gorillas (M. M. Robbins et al., 2022). For all gorilla populations, a substantial proportion of their non-fruit diet consists of protein-rich terrestrial herbaceous vegetation (THV) (Rothman et al., 2006). Within gorilla species, patterns of home range size relate to fruit and THV availability. For instance, in western gorillas, monthly home range sizes increase with a higher percentage of fruit in the diet (Seiler & Robbins, 2020), and in mountain gorillas, monthly home range sizes decrease as the availability of herbaceous foods increases (Seiler et al., 2018). In contrast to gorillas, the diets of both bonobos and chimpanzees are predominantly composed of ripe fruit (70% feeding time in bonobos White, 1992; 75% feeding time in chimpanzees R. W. Wrangham et al., 1998). Within *Pan*, Malenky and Wrangham (Malenky & Wrangham, 1994) argued that bonobos consume more terrestrial herbaceous vegetation (THV) than chimpanzees. Considering only the degree of frugivory, we would predict that home range size of chimpanzees should be larger than that of bonobos, with both *Pan* species having larger home range sizes than both gorilla species (Prediction 1b: C>B>WG>MG). However, in addition to diet, home-range size can reflect numerous interacting ecological variables, including habitat productivity and environmental marginality. For example, among chimpanzees, some of the largest home ranges have been documented in ecologically marginal habitats such as Issa and Fongoli (Piel et al., 2017; Pruetz, 2025), which suggests that habitat quality may be as important as frugivory per se. Adopting this broader ecological perspective, and considering the hypothesized productivity of each species’ environment, we cannot confidently predict differences in home-range size between Pan and Gorilla. We would, however, still predict that chimpanzee home ranges should be larger than those of bonobos, given the comparatively lower productivity hypothesized for chimpanzee habitats relative to bonobo habitats (Prediction 1c: & > B, with no clear prediction for the Pan–Gorilla comparison) (White & Wrangham, 1988).

In addition to total home range size, the economic defendability of a home range depends on the mean daily travel distance, which is broadly influenced by the body size, diet of a given species and the productivity of the environment. Controlling for body size, primate species with a higher reliance on fruits in the diet have to travel longer distances to meet their energetic requirements, because fruits tend to be more patchily distributed in space and time (Clutton-Brock & Harvey, 1977; Milton & May, 1976). Accordingly, travel distance increases with an increase in frugivory in Bwindi mountain gorillas (Seiler et al., 2018). Following expectations based on availability of both fruit and THV, daily travel distances for gorillas are longest in western gorillas (Seiler & Robbins, 2020), and shortest among Virunga mountain gorillas (Wright et al., 2015), with Bwindi mountain gorillas being intermediate (Seiler et al., 2018; Wright et al., 2015). Given the more frugivorous diet of *Pan*, we expect both bonobos and chimpanzees to have longer daily travel distances than gorillas. Bonobos have been proposed to eat more THV than chimpanzees (R. Wrangham, 2023), which would imply shorter daily travel distances for bonobos than chimpanzees, leading to Prediction 2, that travel distances should be ranked as follows: C>B>WG>MG. Environmental productivity has also been linked to daily travel distance across primates, with lower-productivity habitats generally associated with longer travel distances to meet energetic needs (Isbell, 1991; Nunn & Barton, 2000; Chapman et al., 1995). Within apes, similar patterns have been documented, with fruit availability predicting variation in daily travel distance in both gorillas (Doran-Sheehy et al., 2004; Ganas & Robbins, 2005) and chimpanzees (Wrangham et al., 1998). Adopting a broader ecological perspective, differences in daily travel distance between Pan and Gorilla become difficult to anticipate. Within Pan, however, chimpanzee daily travel distances should exceed those of bonobos, as chimpanzee habitats are generally considered less productive (White et al., 2013).

Using home range size and daily travel distance to calculate the defendability index (D), Mitani and Rodman (Mitani & Rodman, 1979) found values of D = 0.15 for Virunga mountain gorillas and D = 0.98 for chimpanzees (no data had been published yet for western gorillas and bonobos). If chimpanzees have both the smallest home ranges (Prediction 1a) and longest daily travel distances (Prediction 2), we would expect D to be highest in chimpanzees, with bonobos intermediate and gorillas with the lowest D (Prediction 3: D: C>B>WG/MG). If, however, more frugivorous diets result in larger home ranges (as in Prediction 1b), the expected order of defendability is not obvious.

In this study we compiled detailed space use data from neighboring groups in four populations of the African apes, one for each species (chimpanzees at Gombe, bonobos at Kokolopori, western gorillas at Loango and mountain gorillas at Bwindi) to augment published data from African ape populations living in relatively undisturbed environments. We then tested for the predicted species differences in home range sizes, daily travel distances, economic defendability using all available comparable data and contextualized the ranging data with insights about home range overlap between neighboring groups in the study populations

## METHODS

### Study populations and data collection

For mountain gorillas, we analyzed data from 13 habituated groups in Bwindi Impenetrable National Park, Uganda (Seiler et al., 2018). We restricted the analysis for this study to the 12 consecutive months with highest data availability (July 2012 to June 2013; monthly mean per group =16 days [range: 2–31 days]; SI Table 2). Location data of the groups were collected in 30 second intervals while directly following the groups for 4 hours, which is the observation time allowed by the park authority regulations to limit anthropogenic disturbance. The remainder of the daily travel data were collected by walking along the trail left by a gorilla group on a respective day. These trails are easily detectable by bent vegetation, dung and food remains.

For western gorillas, we analyzed data from Loango National Park, Gabon, from two neighboring groups, Atananga (ATA) and Evoto (EVO), during 2021 and 2022. Travel data were collected either by following the gorillas in view or by following the trails left where the gorillas had walked (assessed by bent vegetation, discarded plant remains, hand and footprints, and dung). While the Atananga group (ATA) was fully habituated during the study, the Evoto group (EVO) was only followed since 2018 and at the time of data collection not fully habituated. Location data were collected on both groups every 30 to 60 seconds using handheld GPS units (GPSmap 60CSx and 62), resulting in 692 days of data (ATA: N_2021_ = 249; N_2022_ = 246; EVO: N_2021_ = 97; N_2022_ = 100) (SI Table 2).

For chimpanzees, we analyzed data from the long-term study at Gombe National Park, Tanzania. We examine data from three chimpanzee communities: Mitumba (MT) in the far north (observed 1985-present), Kasekela (KK) in the center (observed 1960-present), and Kahama (HK) further south. HK separated from KK by 1973 (Feldblum et al., 2018) and was observed from 1973 to its extinction in 1977 (Goodall, 1986). We included data from KK from 1973 (first year of systematic focal follow data throughout their range) to 2016 (most recent year for available data in database; N_KK_ = 44 years; total number of follows = 12,072). For MT, we included data from 1997 (start of reliable home range estimates (Mjungu, 2010)) to 2016 (N_MT_ = 20 years; total number of follows = 4,580). We also included the sparse ranging data from HK in 1973 (N_1973_ = 31) and 1974 (N_1974_= 61). From 1973 through 2015, observers recorded the focal target’s location at 15-minute intervals on paper maps, which in the 1990s were redrawn from aerial photos rectified using Global Positioning System (GPS) measurements (Gilby et al., 2006). Starting in 2016, observers used hand-held GPS units, which recorded the observer’s location throughout each follow.

For bonobos, we analyzed data from three neighboring communities in the Kokolopori Bonobo Reserve, Democratic Republic of Congo. A team of researchers followed focal parties (staying with one subgroup throughout the day) of the Ekalakala (EKK), Kokoalongo (KKL) and Fekako (FKK) communities from January 2021 to December 2022. When following bonobos, observers recorded their location every 60 seconds using handheld global positioning system units (GPSmap 60CSx and 62), resulting in a total of 1066 observation days with location data (EKK: N_2021_= 193; N_2022_ = 243; KKL: N_2021_= 200; N_2022_ = 231; FKK: N_2021_ = 101; N_2022_= 98) (SI Table 2).

### Home range size

We fit 95% kernel density estimates of each group’s yearly home range based on the location data available for each species (Worton, 1989). This includes the GPS locations from the group follows of western and mountain gorillas, the focal follows of chimpanzees (regardless of the duration), and the party follows of bonobos. We characterized the size of a home range as the area under 95% of the utilization distribution, or the area for which the cumulative probability of use by that group is equal to 0.95. We used the kernelUD function with a bivariate kernel from the R package adehabitatHR (v. 0.4.22) to fit the utilization distribution and calculated area by bounding the utilization distribution at 95% using the getverticeshr function (Calenge, 2024).

To compare our measures of home range size with data from other groups of African apes, we extracted this information from publications (SI Table 3). For home ranges of chimpanzees and bonobos we only considered publications that included 95% kernel density area estimates (SI Table 3). To ensure ecological comparability, we only included chimpanzee data from groups that were not heavily disturbed by human activities. If there were multiple values for a given group, we prioritized measures of annual home range sizes over cumulative values (see overview of incorporated published datasets in SI). Much of the published gorilla data used 90% rather than 95% kernel density estimates. To compare our results with this published data, we calculated both 95% and 90% Kernel density estimates with our gorilla data from Bwindi and Loango and then calculated a translation factor (1.27; Range 1.17-1.39, SI Table 4) that we multiplied with the published 90% kernel data in order to include these studies in the larger dataset (SI Table 3/4).

### Daily travel distances

#### Mountain Gorillas

We used the daily travel distances from the published data set of (Seiler et al., 2018). In this dataset the daily tracklogs were already smoothed, including only GPS points with a minimum distance of 25m to previous chosen points and further reducing the area between the newly chosen points and the track defined by the already chosen points (Seiler et al., 2018).

#### Western Gorillas

We calculated daily travel distances on the tracklogs from the group follows. Given the presence of elephants in the forest and the associated risk of walking in the forest in the dark, these tracklogs did not always start and end at the exact nest site of the gorillas. We included all tracklogs that had at least 8 hours confirmed contact. Due to GPS measurement error (commonly 15–20 m) and a consequential potential overestimation of the path length, we smoothed the tracklogs of the focal follows by filtering GPS recordings to ensure a minimum of 40m distance between consecutive points. We only included daily travel distances from the ATA group (N_2021_=162; N_2022_=128); the EVO group was not fully habituated and so human observation could have affected travel distance.

#### Chimpanzees

We calculated daily travel distance for each follow based on the summed distance between successive map locations (Gilby et al., 2013) or directly from GPS logs (from 2016). We restricted analysis to a subset of follows for which the observers 1) recorded ≥12 hours of observation and 2) followed the target from their morning nest to their night nest (N_KK_ = 3,564 follows; N_MT_ = 715 follows; N _HK_ = 13 follows). We calculated daily travel distances separately for males and females.

#### Bonobos

In addition to the party follows, observers followed individual bonobos from the time the focal bonobo left its nest in the morning until it constructed its nest in the evening and collected their location data every 60 seconds by the same means as the focal party location data. Given the very dense environment, the focal individuals are often out of sight, even if they are part of the focal party. We calculated daily travel distances for individuals with uninterrupted full day focal follows (from nest in the morning to nest in the evening), for which we had at least 8 hours direct observation (N_EKK_=101; N_KKL_=79; N_FKK_=41). Due to GPS measurement error, we applied the same smoothing procedure as the western gorilla data. We calculated daily travel distances separately for males and females.

To compare our measures of daily travel distance with data from other groups of African apes, we extracted information of travel length from publications (see overview of incorporated published datasets in SI; SI Table 3).

### Defendability of the home range

We calculated a defendability index for home range sizes of all groups of African apes following the method laid out in Mitani and Rodman (Mitani & Rodman, 1979). The index of defendability (D) is the ratio of average daily travel distance (d) to the diameter of a circle with an area equal to that of the observed home range (A).

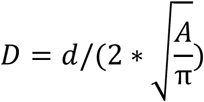

Territorial species are expected to have D >1, indicating they could cross the full width of their home range on an average travel day.

To compare our measures of D with data from other African ape groups, we calculated D for all groups for which both home range size and daily travel distance have been published (overview of incorporated datasets in SI Table 3). Given unavoidable differences in methodology, we derived a comparability index to indicate how directly comparable the estimates are. We scored on a binary basis (yes = 1, no = 0) whether each of the following criteria were met: (1) home ranges reported as annual means; (2) daily travel distance based on full-day follows; and (3) estimates for home range size and daily travel distance were based on data from the same years of observation. The comparability score is the sum of these values (0–3).

### Home range overlap

We characterized home range overlap based on all location data available for each species by calculating a probability density measure with the function kerneloverlaphr (method = “PHR”) from adehabitatHR (Calenge, 2024). The value of overlap PHR_i,j_ corresponds to the probability that an animal from group *j* is found in the home range of group *i*. This method takes better advantage of the kernel home range estimates than traditional area overlap metrics and more closely provides information about the likelihood that animals from different groups would encounter each other (Fieberg & Kochanny, 2005).

To characterize the overlap or shared use of space by different groups, we needed all estimates for a year/site to be calculated with the same bandwidth. We therefore estimated the utilization distributions of multiple groups in concert, as it is important to use a sensible smoothing bandwidth that does not over- or under-smooth any particular group at the expense of others (Fleming & Calabrese, 2017). By default, adehabitatHR uses a ‘reference’ bandwidth calculated by assuming that space use is normally distributed that is highly influenced by home range size (Calenge, 2024). When home range sizes and amount of data did not differ much between groups and when there were fewer groups, the jointly calculated reference bandwidth could be used for all groups (the argument same4all = TRUE in the kernelUD function). When this method failed, we used the mean of all groups’ reference bandwidths, which produced sensible estimates (Silverman, 1986; Wand & Jones, 1994).

For mountain gorillas, the 13 groups did not all live in neighboring areas, but rather in three regions, so it was not computationally feasible to calculate all groups’ utilization distributions in concert. To perform the estimates but maintain our ability to calculate overlap, we split the groups into three “regions” of groups with potential overlap as assessed by a visualization of location data (Ru, Mu, and Ha; Ky and Bi; and Bu, Bw, Kah, Kak, and Mi; (Seiler et al., 2017). We then estimated home range area with the mean reference bandwidth per region as described above.

We compared the maximum observed overlap per population as it better demonstrates the potential for home range overlap within a given population than an average measure.

### Data analysis

To test the predictions of the *economic defendability hypothesis*, we analyzed data from all the African apes relating to three key variables: total home range size (Prediction 1a/b), daily travel distance (Prediction 2), index of defendability (Prediction 3) and considered them in the context of the degree of home-range overlap. We initially compared patterns in these metrics across our study populations (SI Table 2), then integrated them with comparable published data (SI Table 3) and performed formal statistical analyses using a series of linear models. For home range overlap, we examined data from our long-term studies. Because few other studies have published directly comparable measures of home range overlap, the small sample size precluded conducting formal statistical tests.

A first model (Model 1) tested for species difference in home range size (based on published values and the ones extracted from our study groups) (Prediction 1a/b/c). We used the area of 95 KDE of home ranges as a response variable and included species as a predictor while controlling for number of adult individuals within groups as a fixed effect. A second model (Model 2) tested for species differences in travel distance (based on published values and the ones extracted from our study groups) by having the daily travel distance as response variable and species as predictor (Prediction 2). Given the uncertainty to which we expected group size to affect daily travel distances in species with fission-fusion dynamic, we ran the model with and without controlling for the number of adult group members as a fixed effect (Model 2a and 2b). Finally, we conducted a third model (Model 3) to assess species differences in D, using the index of home range defendability as the response variable and species as the predictor.

Because published data did not always include both home range size and daily travel distance, and to control for the possibility that variation in methodology across sites could bias estimates of these measures in different directions, we visualized the parameter space for each species with a rectangle defined by the published range of estimates for home range area and daily travel distance, and plotted a line for D =1 across the range of daily travel distances; points above this line are expected to be less economical to defend (Figure 1). While Mitani and Rodman (Mitani & Rodman, 1979) proposed D ≥ 1 as an approximate threshold above which territorial defense becomes economically feasible, subsequent work has questioned the general predictive value of this specific cutoff across taxa (Lowen & Dunbar, 1994a). While we indicate this line visually, we treat D primarily as a continuous, comparative index of relative defendability across species and populations, rather than as a strict categorical threshold.

**Figure 1.**
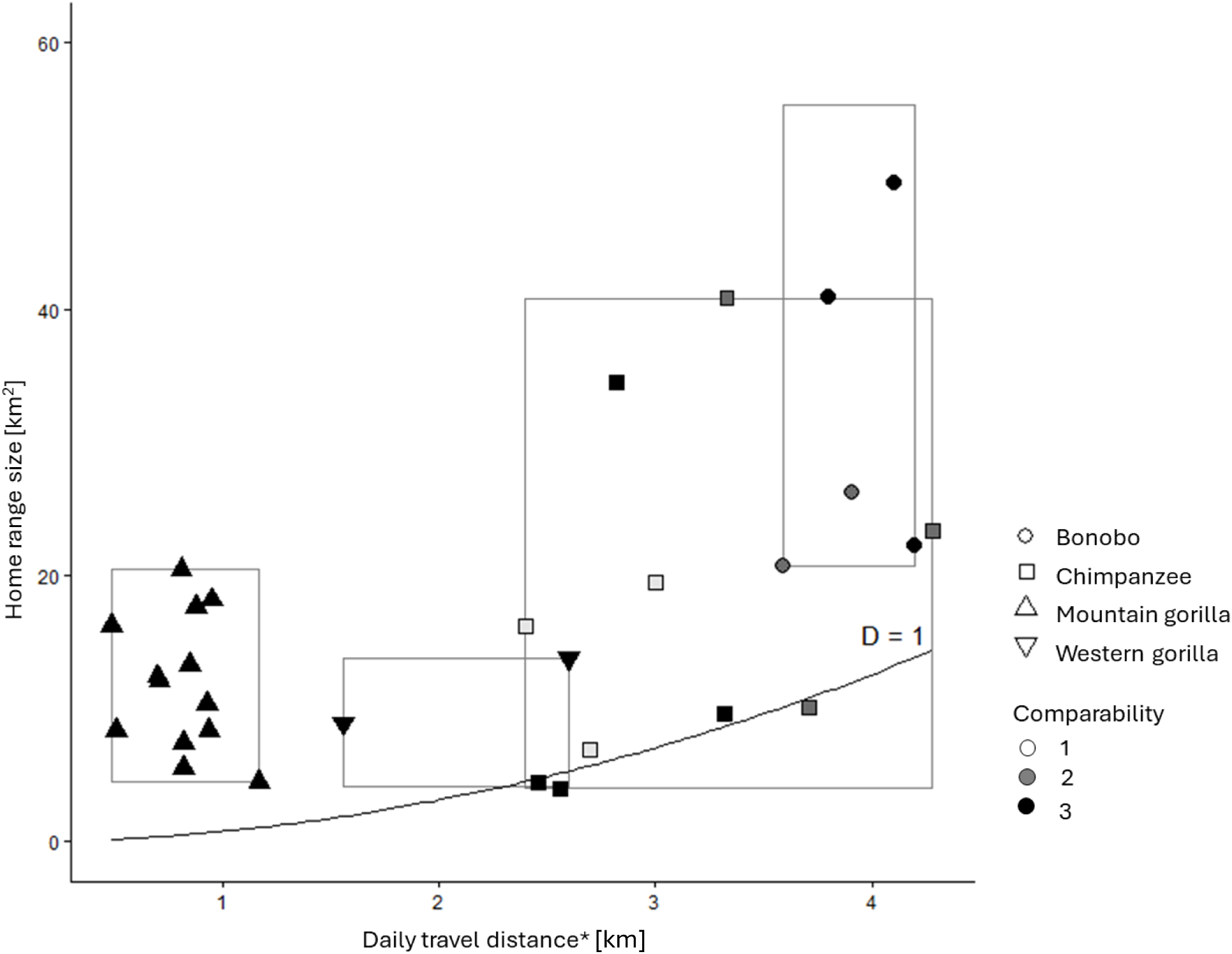
shows the relationship between 95% kernel density home range size [km^2^] and daily travel distance [m] in African apes, combining our data with published values (symbols denote species; see SI for details). Each datapoint represents a group. Boxes indicate broad species-specific parameter spaces, defined by the minimum and maximum annual home range sizes and daily travel distances from our combined dataset. Shading indicates the comparability of each datapoint to the data from this study, based on three criteria: (1) home range reported as an annual estimate; (2) daily travel distance based on full-day focal follows; and (3) overlap in the time periods for the home range and daily travel distance estimates. The comparability score is the sum of the criteria met (0–3). The solid line indicates an economic defendability index D = 1, where daily travel distance equals the diameter of the home range, such that points below this line are within the range in which it is economically feasible to defend the home range according to (Mitani & Rodman, 1979).

Given that in species with fission fusion dynamics, the sexes may differ substantially in daily travel distance with implications for defendability, we tested for sex-differences in daily travel distance in chimpanzees and bonobos.

## RESULTS

### Home range size

The median estimated annual home range area of mountain gorilla groups was 14.00 km2. There was considerable variation between the 12 groups, with estimates ranging from 6.11 to 29.37 km2 (SI Table 2). The median estimate annual home range area of the western gorilla groups at Loango was 17.63 km2. Area estimates ranged from 15.42 to 21.71 km2 across the two groups and years. The annual home range sizes of Gombe chimpanzees varied substantially over time but fell within the range of gorilla home range sizes with a median estimated area of 4.47 km2. Area estimates ranged from 3.02 to 18.42 km2 across groups and years and increased significantly (p-value of a t-test against β = 0 was less than 0.01) year-on-year over the course of the study for both the Kasekela (β = 0.22) and Mitumba (β = 0.15) communities.

The annual home ranges of bonobo communities at Kokolopori were larger than those of the other three species of African apes in our sample with a median estimated area of 38.31 km^2^ (SI Figure 1). Estimates varied considerably between groups (EKK Median = 40.96 km²; KKL Median = 49.43 km²; FKK Median = 22.29 km²; SI Table 2).

Integrating our data into a larger compiled dataset of published data providing 95% Kernel density area estimates (SI Table 3), we found that bonobos had overall significantly larger annual home range sizes compared to similar sized groups of the other African ape species (Figure 1, Table 1, SI Figure 2).

**Table 1.** Results from models investigating ranging patterns based on our and published data in the African apes. Estimated coefficient values for chimpanzees and gorillas are given in relation to bonobos. T- and p-values are reported from t-tests against the hypothesis that each estimate is equal to 0.

| Response variable | Predictor variables | Median of response (Range) | Sample size | Estimate* | SE* | t-value * | p * |
|---|---|---|---|---|---|---|---|
| Home range size [km2] | *Intercept* |  |  | 32.05 | 4.6 | 6.95 |  |
|  | Bonobo | 33.58 (20.7-55.4) | 6 |  |  |  |  |
|  | Chimpanzee | 17.3 (4.01-38) | 15 | -20.84 | 4.99 | -4.18 | <0.001 |
|  | Mountain gorilla | 11.29 (4.35-43.87) | 24 | -19.66 | 4.63 | -4.25 | <0.001 |
|  | Western gorilla | 7.87 (4.17-13.8) | 7 | -23.70 | 5.55 | -4.26 | <0.001 |
|  | *Number of adults* |  |  | 0.24 | 0.15 | 1.63 | 0.11 |
| Daily travel distance [km] | *Intercept* |  |  | 3920 | 200.8 | 19.52 |  |
|  | Bonobo | 3.91 (3.59-4.20 | 5 |  |  |  |  |
|  | Chimpanzee | 2.89 (2.1-4.2) | 10 | -1034.4 | 245.9 | -4.21 | <0.001 |
|  | Chimpanzee males** | 2.91 (2.4-4.28) | 10 | -837.9 | 231.5 | -3.62 | 0.001 |
|  | Mountain gorilla | 0.82 (0.49-1.17) | 13 | -3213.2 | 236.3 | -13.6 | <0.001 |
|  | Western gorilla | 2.08 (1.56-2.6) | 2 | -2158 | 375.7 | -5.74 | <0.001 |
| Economic defendability Index [D] | *Intercept* |  |  | 0.64 | 0.07 | 9.10 |  |
|  | Bonobo | 0.68 (0.52-0.79) | 5 |  |  |  |  |
|  | Chimpanzee | 0.66 (0.43-1.13) | 10 | 0.10 | 0.09 | 1.15 | 0.26 |
|  | Chimpanzee males** | 0.68 (0.43-1.13) | 10 | 0.14 | 0.09 | 1.65 | 0.11 |
|  | Mountain gorilla | 0.55 (0.48-0.62) | 13 | -0.46 | 0.08 | -5.58 | <0.001 |
|  | Western gorilla | 0.2 (0.11-0.49) | 2 | -0.20 | 0.13 | -1.50 | 0.15 |

\* in relation to bonobos \*\* values from separate model including male instead of values for both sexes

### Daily travel distance

To test for species differences in daily travel distance, we compared daily travel distances in our study populations and published results.

Mountain gorillas at Bwindi traveled a median of 707m per day (N=12 groups; range: 108–6,455 m), with substantial variation in annual measures of median daily travel distance among groups (range: 419–1,051 m; SI Table 2, SI Figure 1). Among western gorillas at Loango, the ATA group’s median daily travel distance was 1,527 m in 2021 and 2,400 m in 2022 (SI Table 2). For chimpanzees at Gombe, daily travel distance differed between the sexes for both the KK and MT chimpanzee communities, with males traveling significantly farther (median_KK_= 3323 m, range_KK_: 2059-4881 m, n _KK_=43; median_MT_ = 2457, range_MT_: 1227-3403 m, n _KK_=16) per day than females (Median_KK_=2638m, range_KK_=2070-4125m, n _KK_=43; median_MT_=2331, range_MT_=1147-2869m, n _KK_=14; GLMM controlling for group and ID: χ² = 29.25, df = 1, p < 0.001; see SI Figure 3). However, this difference decreased over time (KK: β = –25.9, p < 0.01; MT: β = – 51.4, p < 0.01). Among bonobos at Kokolopori, daily travel distances were greater than those of the other African apes sites, with little annual variation between communities in median daily travel distance (median_EKK_ = 4,125 m, range_EKK_: 4002-4246m, N_EKK_ = 2 years; median_KKL_ = 3,972 m, range_KKL_: 3957-4128m, N_KKL_ = 2 years; median_FKK_ = 3,923 m, range_EKK_: 3575–4270m, N_FKK_ = 2 years; SI Table 2, SI Figure 2). Additionally, the sexes did not differ significantly in daily travel distance (GLMM controlling for group and ID: χ² = 2.63, df = 1, p = 0.10; see SI Figure 3). Integrating our data into a larger compiled dataset of published data on daily travel distances (SI Table 3), we found that bonobos had overall a significantly larger daily travel distances than the other African ape species (Figure 1, Table 1, SI Figure 1/2). Further, both chimpanzees and western gorillas traveled significantly further during a day than mountain gorillas, and chimpanzees traveled significantly further than western gorillas (SI Figure 2). For all chimpanzee sites for which ranging data from males and females exist, males tend to have longer daily travel distances (SI Table 3).

### Home range defendability and overlap

The economic defendability index D in our study groups of mountain gorillas ranged between 0.09 and 0.33 (N=12 groups), and 0.34 and 0.50 for the one group of western gorillas. This index differed among the chimpanzee communities (Median D_KK_=0.90, range_KK_: 0.58-1.41; median D_MT_ =0.95, range_MT_: 0.58-1.36; median D_HK_=1.11, range_HK_ = 0.82-1.40) and decreased over time for Kasekela (*β* = −0.01, t = −8.35, p<0.01), but not for Mitumba (*β*= −0.02, t=-1.29, p=0.22). D was higher when considering only male daily travel distances (Median_KK_=1.01, range_KK_: 0.55-1.90; median_MT_ =1.06, range_MT_: 0.53-1.54; median_HK_=1.12 range_HK_ = 0.84-1.40). In bonobos, we found considerable variation in D between communities ranging from a yearly minimum of 0.48 in KKL and a yearly maximum of 0.80 in FKK (SI Table 2). Augmenting our dataset with Ds derived from published home range and travel distance data from other African ape populations (SI Table 3), we found that chimpanzees and bonobos had a significantly higher defendability index than mountain gorillas but that no significant differences existed between the species of the genus *Pan* and western gorillas (Figure 1, Table 1, SI Figure 2). Including D values for chimpanzees that are based solely on male daily travel distances does not change the results (Figure 1, Table 1, SI Table 1, SI Figure 4, SI table 4). D values based on daily travel distances of both sexes and of males only for the eastern and western chimpanzee subspecies overlapped completely (SI Figure 5). For chimpanzees, the parameter space of published daily travel distances and home range sizes encompassed a substantial area where D > 1. Of the ten chimpanzee study groups, half (N=5) had defendability indices close to D = 1 (Figure 1). For western gorillas, a small corner of the parameter space encompassed D = 1, but all groups with estimates for both home range size and daily travel distance had points well above this line (D < 1). We note that D = 1, as originally proposed by Mitani and Rodman (1979), represents an approximate heuristic rather than a validated categorical threshold, and we therefore interpret D primarily as a continuous, comparative measure of relative economic defendability across species and populations (REF).

The extent of home range overlaps between neighboring groups of mountain gorillas at Bwindi varied substantially with a maximum probability of 96% that an animal from one group would be found in another group’s home range (SI Table 2, SI Figure 1). Similarly, the home ranges of the western gorilla groups at Loango largely overlapped, with estimates of the same probability measure up to 87% (SI Table 2; SI Figure 1). In chimpanzees at Gombe, home ranges between neighboring groups overlapped very little with a maximum value of 12% probability of presence of Kasekela chimpanzees in Kahama’s home range in 1974 (SI Table 2; SI Figure 1). In the bonobos at Kokolopori, home range overlap between neighboring groups varied substantially but reached as high as a 79.4% probability of presence of FKK in KKL’s home range in 2022 (SI Table 2; SI Figure 1).

## DISCUSSION

Our quantitative analysis of ranging data supports the view that most African ape species do not defend territories because doing so is not economically viable. Contrary to predictions based on the view that bonobos face higher food availability, they had larger home ranges and longer daily travel distances than chimpanzees, implying that they must travel farther to meet their nutritional needs. Economic defendability indices did not differ significantly among chimpanzees, bonobos, and western gorillas, indicating that the feasibility of economic territory defense is not underlying the stark species differences in intergroup behaviour. However, unlike the other apes, roughly half of the chimpanzee groups had indices close to D = 1, which Mitani and Rodman (Mitani & Rodman, 1979) originally proposed as an approximate benchmark for the feasibility of economic defendability.

### Home range size

We found that the sizes of African ape home ranges overlapped to a considerable degree, but when controlling for group sizes, bonobos had the largest home ranges. This contradicts predictions based on body size (Prediction 1a: MG/WG>C/B), diet (Prediction 1b: C>B>WG>MG), and general environmental productivity (Prediction 1c: & > B), but is consistent with the observation by Pusey (Pusey, 2022).

It seems unlikely that our findings are due to biased ecological conditions in the study populations, as the data comes largely from forest-dwelling apes for which comparable data are available. However, published accounts from the Manzano bonobo group, inhabiting a forest–savanna mosaic where movements are likely constrained by habitat fragmentation, indicate a home range size of 20.3 km² by MCP, (group size: 10 adults) (Pennec et al., 2020), which is small for bonobos. Although gorillas do not inhabit such habitats, available data from savanna-dwelling chimpanzees (including the Fongoli population) suggest a different pattern in response to open environments, namely larger, rather than smaller, home ranges, presumably due to reduced resource availability (Lindshield et al., 2021).

Despite the expectation that more frugivorous western gorillas should have larger home ranges than mountain gorillas, our results align with previous work showing that both species have similar home range sizes (Caillaud et al., 2014; Judson et al., 2024; Seiler et al., 2018; Seiler & Robbins, 2020). Our finding that bonobos have larger home range sizes than both mountain and western gorillas is consistent with their greater reliance on fruit and the general pattern across primates whereby fruit-rich diets are associated with larger home ranges.(Clutton-Brock & Harvey, 1977). However, the finding that bonobos also appear to have larger home ranges than chimpanzees contrasts with the pattern expected if bonobos consume less fruit and more on THV than chimpanzees (Malenky & Wrangham, 1994), and also challenges the prevailing view that bonobos benefit from greater food abundance and a higher environmental productivity than chimpanzees (White, 1998).

### Daily travel distance

Our expectation that daily travel distance would correlate closely with proportion of fruit in the diet (Prediction 2a: C>B>WG>MG) received partial support. Western gorillas traveled farther per day than mountain gorillas, as expected given their more frugivorous diet and the lower abundance of THV in their environment (Seiler & Robbins, 2020). Both Pan species also showed significantly greater daily travel distances than gorillas, consistent with their higher reliance on fruit (Clutton-Brock & Harvey, 1977; Koenig, 2002). Contrary to the common assumption that bonobos face lower within-group competition due to a more abundant, stable food supply or reduced dependence on ripe fruit compared to chimpanzees, we found that bonobos actually had longer daily travel distances than chimpanzees and thus appear to travel farther to meet their nutritional needs.

One potential explanation could be that bonobos show higher female gregariousness in their fission fusion dynamic and consequently larger subgroups compared to chimpanzees (Furuichi, 2011; Surbeck et al., 2021). The expected associated increase in competition over food in bonobos could explain their need for increased daily travel distances under ecologically similar conditions. Thus while large daily travel distances are also reported in the Taï chimpanzee population, where females form larger parties than at many other sites (Samuni et al., 2021), the smaller party sizes in many chimpanzee populations may reduce the need for longer travel. This raises the question why bonobo females travel in larger parties rather than minimizing travel distances, which would seem energetically advantageous. Overall, while increased female gregariousness in bonobos may explain their longer travel distances, it does not account for species differences in home range sizes, assuming the depletion of food resources depends mainly on number of individuals feeding in a given area.

Together with generally smaller group sizes, the larger home ranges and travel distances in bonobos compared to chimpanzees suggest an overall reduced abundance of high-quality resource availability and challenge the notion that bonobos experience higher food abundance and/or a more gorilla-like diet relative to chimpanzees (White et al., 2013; R. W. Wrangham, 2000). At this stage, more accurate, comparable measures of food availability across time and space, and of interspecific food competition (Head et al., 2011; Morgan et al., 2016), would greatly improve assessments of ecological differences among African apes. Furthermore, given variation in home range overlap, population density, rather than group size or home range extent, may better capture the nutritional resources available to each species at a site (Jetz et al., 2004; Pearce et al., 2013).

### Defendability and home range overlap

The defendability indices for both gorilla species were well below the heuristic threshold indicative of economic defendability inferred by Mitani and Rodman (Mitani & Rodman, 1979), which is consistent with the absence of territorial behaviour and in line with predictions of the economic defendability hypothesis. Both species in the genus *Pan* exhibited significantly larger defendability indices than mountain gorillas. Bonobos had larger home ranges than chimpanzees, but also traveled further each day, giving them defendability indices similar to those of western gorillas and overlapping with the range of values observed in chimpanzees. Chimpanzees exhibited considerable variation across study sites, with half the study groups having indices close to the heuristic threshold indicative of a feasible economic defendability. Chimpanzees also vary widely in how often and how intensely they engage in intergroup competition with the two groups at the lowest population densities (Fongoli/Mayebe) lacking reports of territorial behaviour.

Given sex differences in the participation in territorial behavior in chimpanzees and potential sex-differences in travel patterns in species with fission fusion dynamic (Massaro et al., 2022; Samuni et al., 2021), we analysed the travel distances and defendability of home range for both sexes. While we do not find significant sex differences in daily travel distance among bonobos, our finding of sex differences in daily travel distance in Gombe chimpanzees mirrors results from other chimpanzee populations (Bates & Byrne, 2009; Bertolani, 2013; Herbinger et al., 2001). These findings in chimpanzees have been interpreted in light of the joint demands of foraging and territorial defense (Bates & Byrne, 2009). However, including defendability indices based only on male daily travel distances for chimpanzees does not change our results.

One limitation of our broad comparison is that published daily travel distances and home range sizes used to calculate the defendability index were not always collected with the same protocols or during overlapping periods (see SI Table 3). Thus, we cannot rule out that typical issues of large comparative datasets influenced our results. However, even when excluding chimpanzee data where home range and travel distances were estimated from non-overlapping periods, the resulting range of D values still encompasses those observed for bonobos.

Lowen and Dunbar (Lowen & Dunbar, 1994b) proposed a modified defendability index for fission–fusion species, suggesting that multiple subgroups traveling independently could enable defense of larger home ranges. However, chimpanzees typically patrol boundaries in parties containing a large proportion of group males (Langergraber et al., 2017; Massaro et al., 2022), likely to avoid numerical disadvantage in intergroup encounters (Wilson et al., 2001). Although simultaneous patrolling by multiple parties could be feasible in very large communities, it has not been reported in chimpanzees.

This distinction notwithstanding, the existence of alternative formulations of D underscores that no single index or threshold value fully captures the ecological and behavioral factors relevant to quantify the feasibility of territorial defense across species. We therefore remain cautious in our interpretation of the absolute values of D, including the conventional D = 1 benchmark, as an approximate and comparative heuristic rather than a precise or universally applicable cutoff.

Mountain gorillas had the greatest home range overlap, with variation among groups and over time ((Seiler et al., 2017; Seiler & Robbins, 2020) Robbins, pers. obs.). Maximum overlap was lower for western gorillas at Loango, somewhat lower for bonobos at Kokolopori, and lowest for chimpanzees at Gombe. Methodological differences prevent direct comparison with other sites, but the 7.5–13.5% territory overlap reported for Taï chimpanzees ((Herbinger et al., 2001)) appears similar. In contrast, neighboring bonobo communities at LuiKotale show much smaller overlap than at Kokolopori ([71). Although the causes of this variation in bonobos remain unclear, our results show that bonobo communities can have large home range overlap under some conditions and support the view that chimpanzees are the most territorial of the nonhuman African apes.

Even though only about half of chimpanzee groups in our study appear to have home ranges which appear economically defendable, they still show territoriality and intergroup hostility even where ranges are too large to defend (Lindshield et al., 2021), implying that factors beyond range defendability drive their behaviour.

#### Differences in male mating strategies and territoriality

An alternative explanation for variation in territorial behaviour and space use among African apes focuses on intragroup relatedness and strength of reproductive skew. Group cooperation associated with territoriality is hypothesized to arise under conditions of low reproductive skew within groups and high genetic differentiation between groups (Bowles, 2006; Vehrencamp, 1983). In both gorilla species, most groups are one-male, so competition is largely between groups rather than within them; even in the ∼40% of multimale mountain gorilla groups, extremely high reproductive skew (about 85% of offspring sired by the alpha) (Bradley et al., 2005; Vigilant et al., 2015) undermines incentives for male cooperation, despite potential kinship. Male gorillas typically emigrate and pursue individualistic strategies, defending the females of a cohesive group whose members themselves influence group composition and often disperse again (Manguette et al., 2020). In chimpanzees and bonobos, male philopatry could promote intragroup relatedness among males and cooperation in range defense (Willems & van Schaik, 2015). However, in bonobos, a higher reproductive skew potentially reduces the incentives for cooperation and thus male territorial defense (Cheng et al., 2025; Mouginot et al., 2023).

While the concept of economic defendability uses a species’ ranging patterns to explain territoriality, territoriality itself likely influences ranging and grouping patterns by constraining individual movements. This raises an important issue of causal interpretation. Because D is calculated directly from home range size and daily travel distance, and both variables can themselves be shaped by territorial behaviour, D is not entirely independent of the phenomenon it is intended to explain. Mitani and Rodman (1979) themselves acknowledged this potential feedback, and our results likely reflect a similar interdependence. In chimpanzees, territorial defense and the risk of severe violence restrict access to food and shape ranging patterns (Herbinger et al., 2001; Williams et al., 2002). Data from Gombe and other sites show that when neighbourhood pressure declines, for example, after increased mortality in adjacent communities, communities can expand their territories (Mitani et al., 2010; Nishida et al., 1985; Rudicell et al., 2010). Because such territorial expansion directly increases home range size, it would also be expected to alter D itself, illustrating how territoriality can shape the very ranging variables used to calculate defendability, rather than being solely a downstream consequence of them. The resulting increase in territory size, and the possibility of accessing food resources over a larger area, is associated with higher body weight (Pusey et al., 2005) and faster female reproduction (Williams et al., 2004; Wood et al., 2025). Restrictions on movement due to territoriality may also influence fission–fusion dynamics, leading to stronger decreases in party size during periods of low food availability. Consequently, both territoriality and habitat characteristics could help explain why chimpanzees are more sensitive to within-range fluctuations in food availability than less territorial bonobos, who may access areas beyond the group range during scarcity (Lucchesi, Cheng, Deschner, et al., 2020; Surbeck et al., 2021).

Overall, while economic defendability is unlikely to fully account for the observed species differences in territorial behaviour, male competition and mating strategies within groups may contribute to social dynamics between neighbouring groups with cascading effects on other aspects of within-group behaviour, including patterns of space use and association.

## ACKNOWLEDGEMENTS

We thank the governments and park authorities of the Democratic Republic of the Congo, Tanzania, Gabon, and Uganda for permission to conduct research: the Ministry of Scientific Research and Technology and the Institut Congolais pour la Conservation de la Nature (Kokolopori); the Tanzania Commission for Science and Technology, Tanzania National Parks, and the Tanzania Wildlife Research Institute (Gombe); the Agence Nationale des Parcs Nationaux and the Centre National de la Recherche Scientifique et Technologique (Loango National Park, Gabon); and the Uganda Wildlife Authority and the Uganda National Council for Science and Technology (Bwindi Impenetrable National Park, Uganda), who also supported data collection. We are grateful to the Bonobo Conservation Initiative, Vie Sauvage, and the local communities of Kokolopori for their support and for protecting the reserve; to the Jane Goodall Institute (Gombe Stream Research Centre); and to the Institute of Tropical Forest Conservation and the Uganda Wildlife Authority for logistical support in Uganda. We thank the field staff, assistants, and trackers at all sites for their dedication and long-term data collection. We also thank the many people who have contributed in various capacities to the long-term gorilla datasets in Uganda and Gabon; a comprehensive list is available at https://www.eva.mpg.de/primate-behavior-and-evolution/research-groups/gorilla-group/.

Work at Kokolopori was supported by Harvard University, the Max Planck Society, and the Daughters for Earth and Wild Lives Foundation. Long-term research at Gombe has been supported by many sources, including the National Institutes of Health (R00 HD057992, R01 AI 091595, R01 AI050529, R01 AI120810), the National Science Foundation (DBS-9021946, SBR-9319909, BCS-0452315, IOS-1052693, IOS-1457260, BCS-0648481, BCS-1753437, BCS-1743506), the Arcus Foundation, Carnegie Corporation, the Leo S. Guthman Foundation, Margo Marsh, Mazuri, the Morris Animal Foundation, the National Geographic Society, the Harris Steel Group, the Waitt Foundation, the William T. Grant Foundation, the Windibrow Foundation, and the Jane Goodall Institute. Data collection in Bwindi and Loango was funded by the Max Planck Society, the United States Fish and Wildlife Service Great Ape Fund, Berggorilla & Regenwald Direkthilfe, Tusk Trust, Heidelberg Zoo, and private donors. Open access funding was provided by the Max Planck Society.

